# Measuring Transcranial Magnetic Stimulation-Induced Electric Fields in Anatomically and Conductively Accurate Rat Head Phantoms

**DOI:** 10.1101/2025.10.13.681990

**Authors:** Wesley Lohr, Mohannad Tashli, Chunling Dong, Padmavathi Sundaram, Ravi L. Hadimani

## Abstract

The efficacy of neuromodulation techniques like transcranial magnetic stimulation (TMS), are highly dependent on the geometry and conductivity of the stimulated target of the brain. Although anatomically accurate head models are routinely used for computational simulations of TMS-induced electric (E) felds, there is still a need for realistic phantoms that mimic the anatomy and electrical conductivity of the head. Here, we present a realistic rat brain phantom constructed using advanced 3-d printing technologies. Our phantom allows validation of TMS-induced E-fields using embedded mutually orthogonal triaxial dipole probes (TDP) that can measure induced E-fields along three axes. We tested the TDP probes in the constructed phantom using four TMS coils with different core materials and core geometry. These measurements were then compared to computational simulations using the finite element method (FEM). The rat brain phantoms had a conductivity of roughly 0.5 S/m, which was mimicked in the FEM simulations. When the measured induced e-fields in the phantoms were compared to the simulated results, the measured results were in the same expected range and fairly close to one another with an average error of 5.1%. The peak E-fields (measured vs. simulated) close to the surface of the grey matter were: permender v-tip core (115.3 V/m – 110V/m), permender flat tip core (91.9V/m – 85 V/m), AISI 1010 v-tip core (94.7 V/m – 100 V/m), and AISI 1010 flat-tip core (85.9 V/m – 84 V/m) respectively.

## 1. Introduction

Transcranial magnetic stimulation (TMS) has emerged as a non-invasive technique for stimulating and modulating focal regions of the brain safely. It has been reported to show beneficial effect for many neuropsychiatric disorders that are not currently FDA approved [1-7]. Currently it is an approved treatment for patients suffering from depression, obsessive compulsive disorder, migraines, and smoking addiction [8-11]. Researchers continue to investigate TMS-based treatments for post-traumatic stress disorder (PTSD), traumatic brain injury (TBI), schizophrenia, and anxiety [12-17]. While subcortical regions of the brain are explored as targets for new TMS treatment procedures, existing treatments primarily target superficial cortical areas since TMS shows highest efficacy closer to the surface of the brain, as the magnetic field decays rapidly with the distance from the source of the field [18].

Field penetration depth, field focality, brain geometry, and stimulation parameters such as frequency and intensity are all key variables that dictate the type and efficacy of treatment [18-20]. However, studying and refining these parameters in living subjects, whether human or animal, presents ethical and logistical challenges. In animal studies, euthanization is often the end point of the experiment [21, 22]. Combined with the arduous process of getting Institutional Animal Care and Use Committee (IACUC) approval for animal research, these types of studies can be time consuming and resource intensive. As new treatment modalities are explored and the range of disorders that are FDA approved expands, there is a larger need for TMS phantoms that can replace animal models, and be used to validate the induced E-fields, for those developing new coils and protocols [23].

Many medical imaging modalities including magnetic resonance imaging (MRI), computed tomography (CT) and ultrasound use phantoms for training and device calibration; to the best of our knowledge, such phantoms have not yet been developed for TMS studies. A useful TMS phantom should represent the anatomy and conductivity of the brain and allow for measurement of the induced E-field inside the brain. These phantoms would facilitate validation of TMS protocols, development of new stimulation techniques, and assessment of device performance, before moving on to *in vivo* studies [24, 25]. Here, we present a physically-accurate rat head phantom that allows measurement of the TMS-induced E-field strengths *in-situ*. The 3D printing based approach allowed us to recreate complexities in anatomy that other techniques may struggle with. We used publicly available MRI and CT data [26], along with the SIGMA rat brain atlas [27] to construct an anatomically accurate model using inkjet printing and injection molding. The development of the phantom involved integration of computational image processing of the MRI/CT data with advanced 3D printing techniques. We show how this approach can produce subject-specific brain phantoms. In addition to anatomical accuracy, the technique presented in this paper allows for tunable composite conductivity to match that of brain tissue in engineered phantoms. Furthermore, we have investigated the use of 2 different ferromagnetic cores in the TMS coils and 2 different shapes of cores in combination with firgure-of-eight coils. The choice of ferrmagnatic material and the shape of coil tip was based on our prior publications[28, 29]. While the specific example highlighted here is the rat brain, we expect that the presented approach may also work well for human studies to validate the TMS-induced E-field for different TMS coils.

## 2. Methods

### 2.1 Mesh Construction

MRI and CT images were obtained from public databases to create anatomically accurate brain models. The images were segmented using statistical parametric mapping (SPM) and the SIGMA brain atlases [30, 31]. We preprocessed the data to correct for artifacts, normalized the images to Montreal Neurological Institute (MNI) standard anatomical space, and enhanced the signal-to-noise ratio. We used the MATLAB (Mathworks Inc., Natick, MA) plugin SPM12 to to segment the brain and head regions using binary rat masks from the Sigma rat brain atlas [32-36]. We used 3D Slicer [37] to repair artifacts, resolve overlapping geometries, and convert the segmentations to STL triangle meshes. There was no mask used to create the skin STL since it is easily created in 3D slicer using thresholding. The programs Meshmixer and Meshlab were used to reduce the file size and fix any small mesh errors. The skin, CSF, skull, gray matter, and white matter compartments of the rat’s head were imported into SolidWorks to create 3D-printed molds and parts. Boolean operations were performed to ensure no overlap between the compartments to avoid issues with the simulations and the 3D printing steps. For phantom fabrication, a boolean subtraction between a solid block and the homogenized brain was performed to create a mold of the brain region.

### 2.2 Conductive composite

Polydimethylsiloxane (PDMS) and carbon nanotubes (CNT) were used to create material with the required conductivity. The carbon nanotubes formed a conductive network at the nanoscale level within the polymer. The connectivity of this network was directly related to the bulk conductivity of the material and was simply modified by altering the loading percent of the CNT.

We titrated conductive multiwalled CNTs 20-30 µm long and 80 nm wide (Cheaptubes: Graton, VT) into a PDMS elastomer to assess the range of conductivities for CNT weight percentages ranging from 8% to 15%. Even dispersion of the CNT was necessary for optimal conductivity, so high shear forces and lengthy mixing times were used. A Caframo High-Torque Brushless Mixer was used for up to 10 minutes at 1000 rpm to disperse the CNT. Extreme viscosity changes occur if CNT exceeds 8%. In the mixing stage the composite started with a viscosity similar to honey and ended up as a thick paste, going through three visually distinct viscous phases; the first was the lowest viscosity where it flows and mixes freely; the next was medium viscosity where the composite will begin to climb up the mixing rod and must be continuously pushed back down; in the third phase the composite begins to cling to itself and the walls of the beaker, it no longer flows freely, and we saw a noticeable shift in the ease of mixing. This change in viscosity was the best indicator of even dispersion because the time taken for one batch to reach this viscosity differs. Once the highly viscous stage was reached, the PDMS curing agent was thoroughly mixed into the composite to ensure even curing.

The conductivity of the composite was measured using a Keithley 2400 source meter on 1 cm x 1 cm samples held in a 3D-printed housing. Two metal plates on either side of the sample were connected to the source meter. To ensure good contact between the metal and the samples, a thin layer of conductive carbon paste (MG chemicals) was used to interface these surfaces. The 3D-printed housing ensured that the sample was not compressed and that all samples were of the same dimension. The average conductivity of 8 samples was then taken and plotted for each point.

### 2.3 Triaxial dipole probes

Three triaxial dipole probes (TDP) were embedded in the phantom to measure the E-field components along three mutually orthogonal axes to characterize the E-field vector at a given point. When fabricating the TDPs, we noted the distance between dipole ends (3 mm in Figure. 2) and made sure that it was much smaller than the length of the induced E-field in that plane [39].

**Figure 1.**
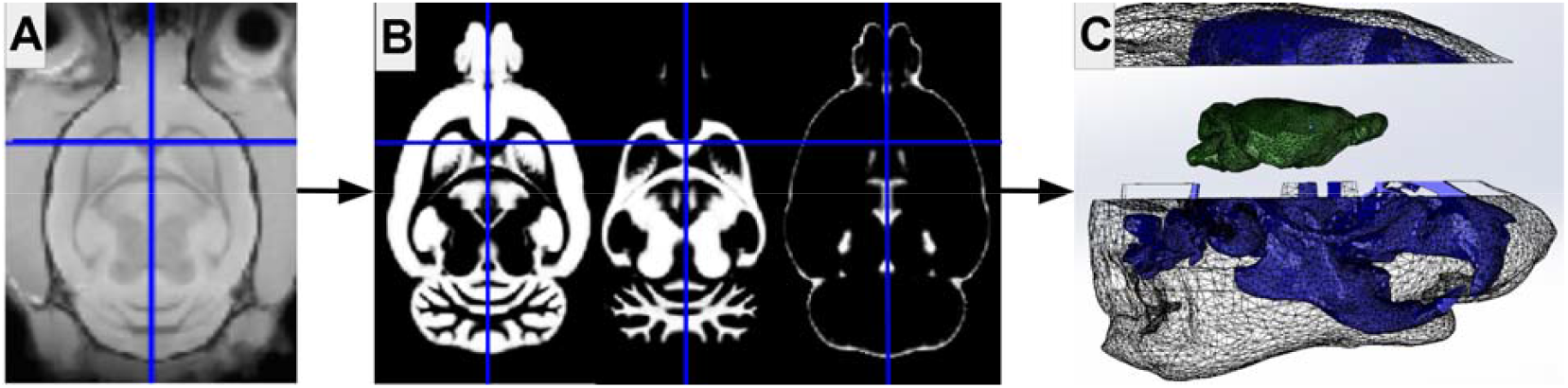
Segmentation of MRI and CT data to construct STL meshes. (A) Co-registered MRI data of the rat head. (B) Segmented grey matter, white matter and cerebrospinal fluid compartments of the rat brain using SPM, thresholding techniques were used to create the skin and skull files. (The blue crosshairs indicate the co-registered origin that all files must share, here the origin is specific to the rat corpus callosum) [38]. (C) Reconstructed STL mesh of the rat head and brain; the skin and skull were 3D printed using a Stratysis J850 polyjet printer; a negative mask of the brain was created and 3D-printed using fused deposition modeling to create a mold for injection molding.

**Figure 2.**
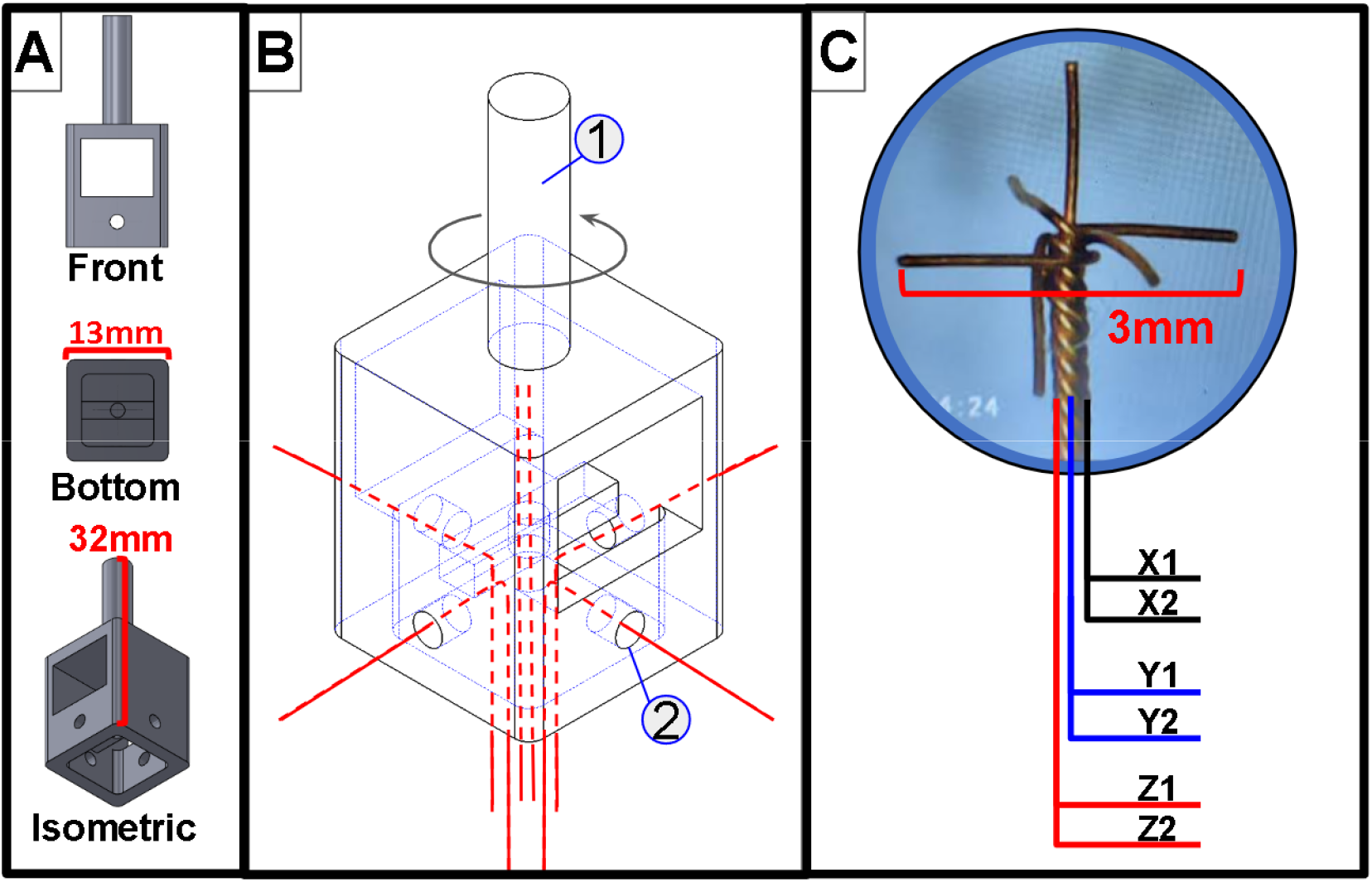
(A) 3D printed TDP winding tool. (B) Six insulated wires were inserted into the 3D printed tool (2), each corresponding to the positive or negative side of a single axis. The tool was then secured into an electric drill using the shaft labelled (1) and the dipoles were wound together. (C) Once wound, the negative Z lead was bent down and the ends were precisely cut to 1.5 mm from the origin. (D) The final result is a TDP with precise axial geometry with known outputs for each axis that are insulated along the length and exposed at the cross section of the dipole’s end.

Six lengths of 37 gauge insulated copper wire were cut at 10 cm. Copper was used because it is less ductile than more conductive metals like gold, or silver and therefore could more easily maintain orthogonality between the different planar dipoles [40]. A 3D-printed housing was designed to wind the TDP while keeping track of the ends of each length of wire. The sides of the housing were labeled as x and y to ensure we knew the axial position of each wire after they were wound together. Point 1 in the **Fig.2** shows a shaft that allowed the housing to be gripped by an electric hand drill. Two housings were used, one at either end, each of the lengths of wire were inserted into the holes labeled as point 2 in **Fig.2** and secured before the drill was activated. Once tight, the wires were cut so that there was 3 mm between the respective dipole ends. This process results in a reliable and repeatable angle between all the dipoles in the xy plane. As for the dipole in the z plane, both ends were held straight up during the process, once the wires were all cut and the TDP was free of the housing, the negative z dipole end was bent down 180 degrees. Two TDPs were placed in each phantom, though the posterior TDPs were not used due to connectivity and placement errors. The TDPs were inserted into the phantoms before the conductive composite was cured.

### 2.4 Polymer injection molding and inkjet 3D printing

The STL mesh models of the brain skin and skull were used to create the phantom. The grey matter and white matter were simplified to a single homogenous brain compartment for ease of creation and replication. A mold was created in SolidWorks by subtracting the brain geometry from a 3 cm x 1.5 cm block and adding ventilation holes and injection ports. TDP were generously coated in the conductive composite then fixed in the proper position in the mold. The conductive composite was loaded into a syringe and injected into the 3D printed mold **Fig. 3a**.The skin and skull were 3D-printed on a Stratasys J850 which allows for a full range of color and flexibility within one part. The skin was flexible and transparent, while the skull was rigid and white. The TDP probes were fed through holes in the bottom of the skull that lead out through the neck and the brain phantom was positioned inside the skull. The top and bottom of the head were then secured together and the TDP leads were soldered to pins in the back of the neck.

**Figure 3.**
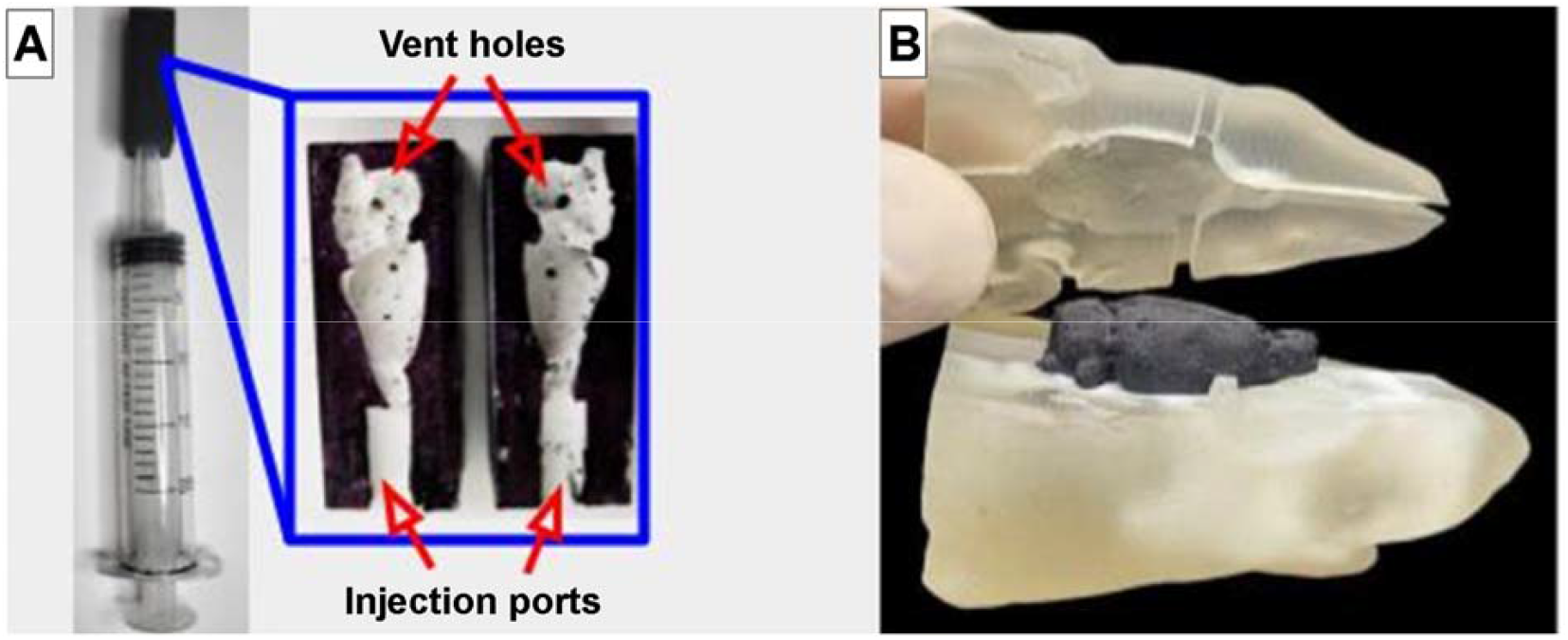
(A) Injection molding of the conductive brain via the use of a disposable syringe and a 3D printed mold with ventilation holes in the 3D printed mold of the rat brain, and injection port to allow flow of the conductive composite. (B) Rat head phantom showing the interior of the skin and skull where the brain rests. A saline solution will be injected into this space under vacuum once the phantom is sealed.

### 2.5 TMS coils

Four figure-of-eight coils like the one seen in **Fig.4** were fabricated with inductance between 10.2 and 15.9 µH, within the inductance range required by the Magventure stimulator [41, 42]. Each cylindrical core had one of two geometries, v tip or flat tip, and one of two core materials, iron-cobalt-vanadium alloy (commercially known as permender) or AISI 1010 carbon steel. The core materials were chosen for their high saturation magnetization, high relative permeability and low core losses [43, 44]. The cores were machined in-house from rods purchased from EFINA. They were then placed in a 3D-printed mount which secures them at exactly the same distance for each coil. The copper windings were made of 3 layers of AWG 26 wire with 8 windings per layer. Once wound, a potting compound epoxy from MG Chemicals was used to secure the windings and insulate the coil. This compound was mixed at room temperature and left to cure for 24 hours before handeling. The change in magnetic field over time is then measured using three orthoginal coil probe.

### 2.6 Simulations

Sim4Life finite element software (Zurich Med Tech, v6.2.1.4972) was utilized to compute the peak intensities of E-fields at 3 planes in the horizontal direction as shown in **Fig.5** within the rat head model, aligning with the experimental E-field TDP measurements. The coils were positioned on top of the rat head model, aligned with the center of the gray matter, and oriented perpendicular to the constructed horizontal planes, aproximately in the same position as in the phantom measurements. The material properties for the segments of the rat’s head—skin, skull, gray matter, white matter, and cerebrospinal fluid—were selected from the IT’IS LF database (IT’IS Foundation, v4.0) [45]. The skin and skull layers have been made invisible in **Fig.5** to more easily visualize the planes of interest in our simulations.

A static vector potential simulation was first conducted to model the coil and account for the relative permeability of the ferromagnetic material. We performed 12 simulations, with 3 for each coil at current intensities of 1000, 666, and 333 Amps, corresponding to 15%, 10%, and 5% of the maximum stimulator output respectively. The results from these coil and core simulations were then used as a source for a magneto quasi-static simulation to explore the effect of the magnetic field on the rat head model.

### 2.7 E-field measurement setup

**Fig. 4** shows the experimental setup of the E-field measurement. The signal originated from the computer, connecting to the DAQ, which then sent the signal to the stimulator then to the coil. For measuring the induced E-field, a dipole probe inserted into our rat head models collected the signal, directing it to the amplifier with a high-impedance head stage which had appropriate filter settings; the upper limit should be sufficiently high to avoid filtering the TMS waveform. The initial ramp-up took approximately 5 microseconds.

**Figure 4:**
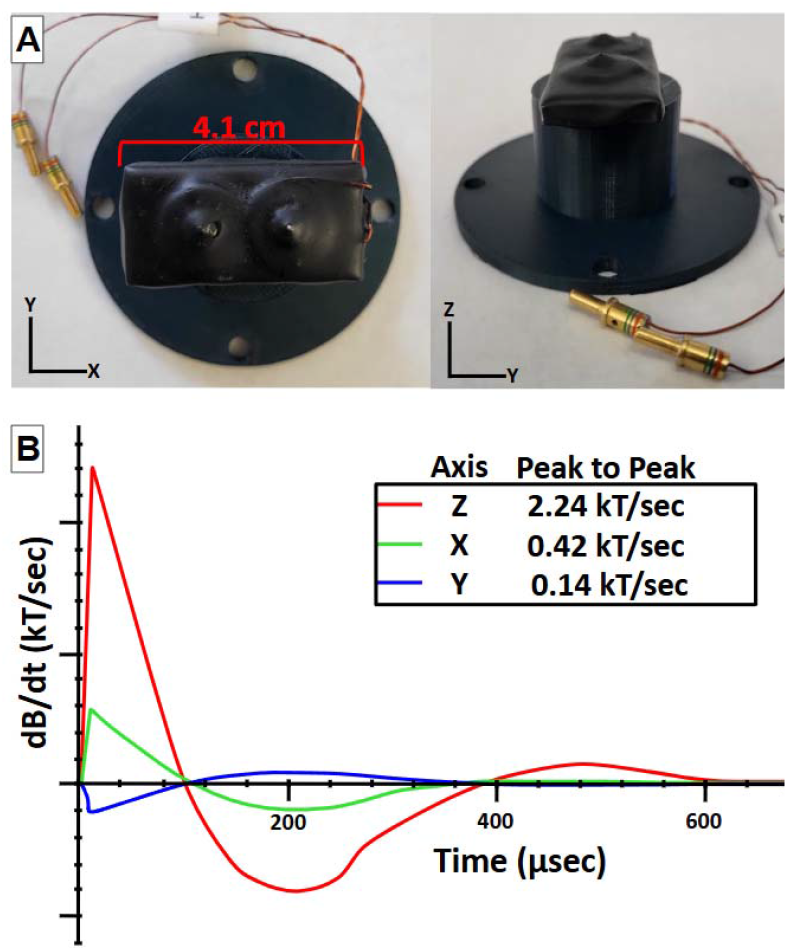
A) Top and side view of the figure of eight coil with a permender v-tip core. The two leads connect to the stimulator. Three more coils with the same number of coil windings and winding layers were also used. B) Biphasic magnetic field waveform of the permender v-tip coil where 1V=1.4 kT/sec.

**Figure 5:**
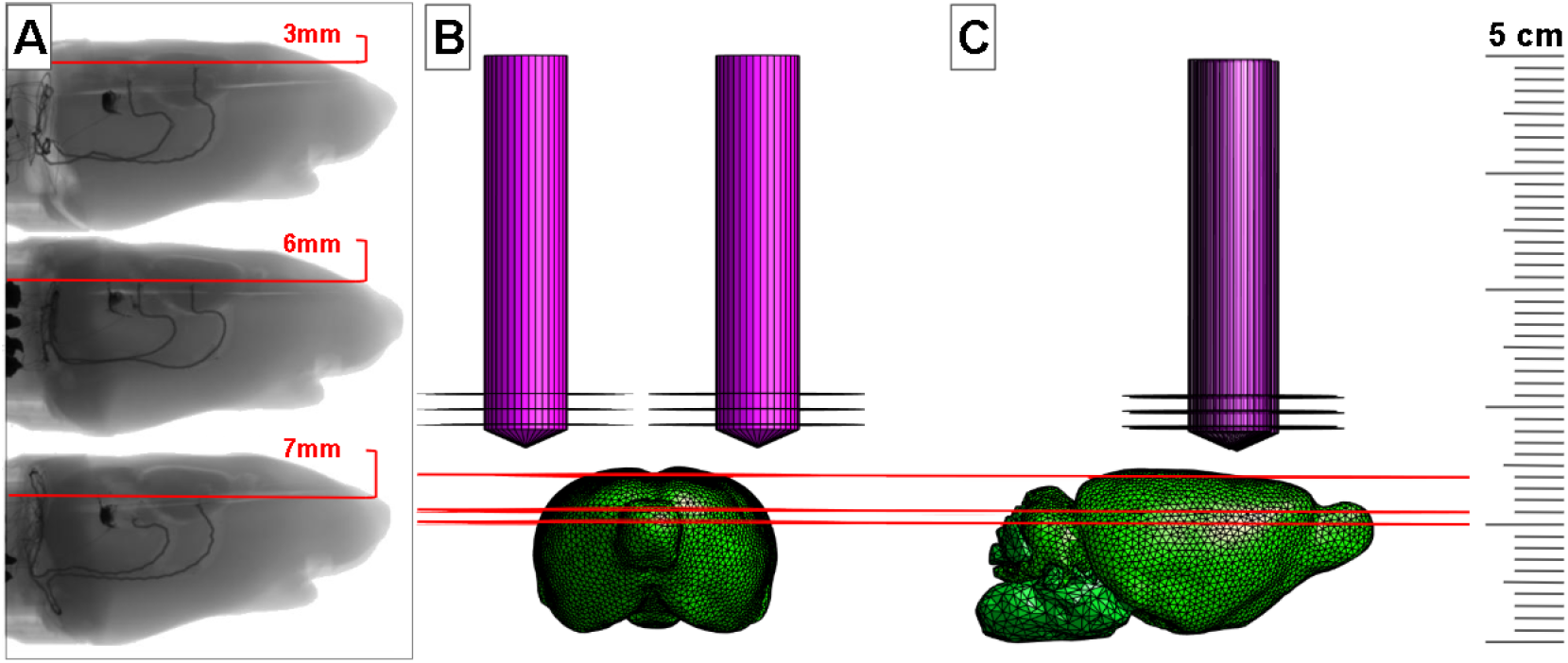
(A) CT scans of the three phantoms used in the experiment. Each phantom has a two TDP probes, the anterior probe captures the e-field signal, while the posterior probe was meant to further characterize the field. (B) Coronal view of the GM STL and coil used for simulations. (C) Sagittal view of the GM STL and coil used for simulations.

Three phantoms (like in **Fig.6)** each had TDPs inserted at different depths. The ends of the dipoles not imbedded in the phantom were soldered to female pins and housed in a 3D printed casing that was fixed to the base of the rat phantom neck in **Fig.6**. This housing allowed the phantoms to be connected to the male pins of what we designate as the TDP switchboard, creating a setup where each phantom was secured in the exact same position relative to the figure of eight coil. Instrumentation limitations mandated that we measure each dipole-end in reference to the same ground point. Three measurements were taken for each dipole end in each phantom, for each of 3 stimulation strengths. This resulted in 6 measurements per dipole, per stimulation strength. The average of these points was taken using to account for variation between the three measurements of each dipole end. Phantoms a, b, and, c were subsequently connected to the switchboard and all measurements were recorded at stimulation strenghs 5%, 10%, and 15%.

**Figure 6.**
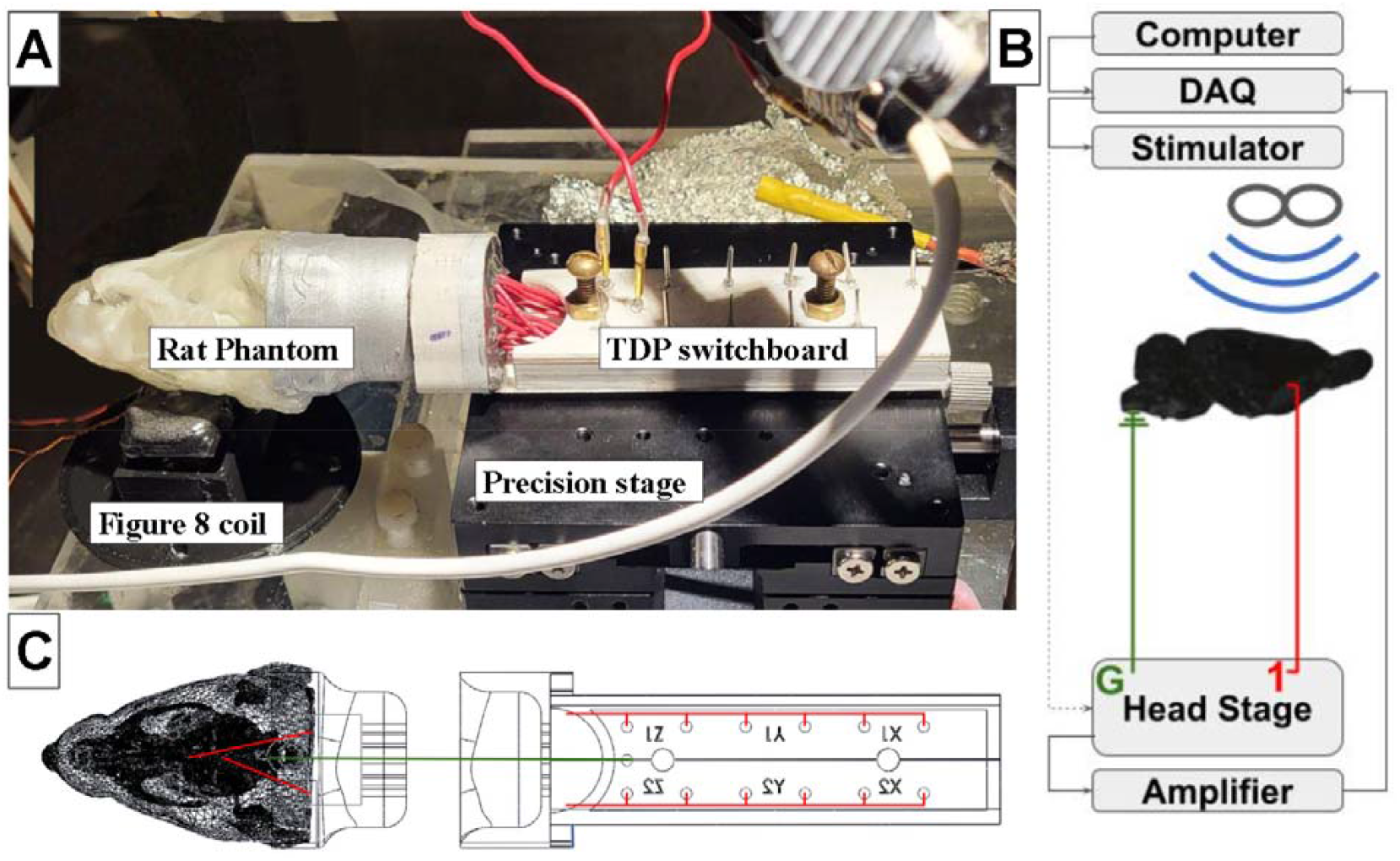
Electric field measurement setup in phantoms. (A) Experimental setup used in measuring e-fields in the phantom, the red wires connect the switchboard (point of measurement on the TDP) to the headstage (not depicted here). (B) Flow chart of the instrumentation used in collecting the e-field signal, depicts ground and the active measuring site on the TDP (C) Schematic of rat phantom and the “switchboard”. Each dipole is measured independently in reference to ground, so an average over multiple pulses was necessary ti capture all components of the e-field at the point of measurement. This board was used to easily switch between dipole measurements.

## 3. Results

### 3.1 Individualized Head Models

The SPM and thresholding techniques used to create the 3D model resulted in an STL mesh file that was representative of the original image data. When comparing **Fig *7*C** to **Fig*7*A-B** some differences can be noted. Most notably, the grey matter and white matter of the brain were combined to one homogenous brain layer, and the trabecular bone in some regions was simplified. The overall geometry of the brain, skin, and skull were preserved in the 3D model, which was used to create the phantoms, and simulated rat head. We took care in creating these models to minimize potentially confounding geometric errors.

**Figure 7:**
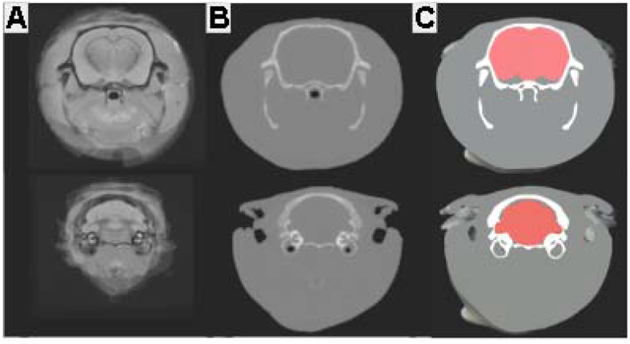
<u>Geometric validation of STL files.</u> (A) MRI slices of Long Evans rat. (B) Slices of CT scan of Long Evans rat. (C) STL slices of Long Evans rat

### 3.2 Conductive composite

The conductivity of the composite was accessed to show the percolation point and the upper limit of the conductive range. The conductivity of the rat brain was averaged to be 0.57 S/m [46, 47], which is why a weight percent of 10.35% CNT was used to create the final composite. In **Fig.8** the brighter regions of the composite represent aglomerations where higher loading percent CNT form mound like structures. At higher weight percentages, these aglomerations would make up most of the surface, leaving it rough and clearly showing the connected network between the CNT. If not properly dispersed using high shear forces, the CNT will not fully infiltrate with the PDMS and form a conductive network.

**Figure 8:**
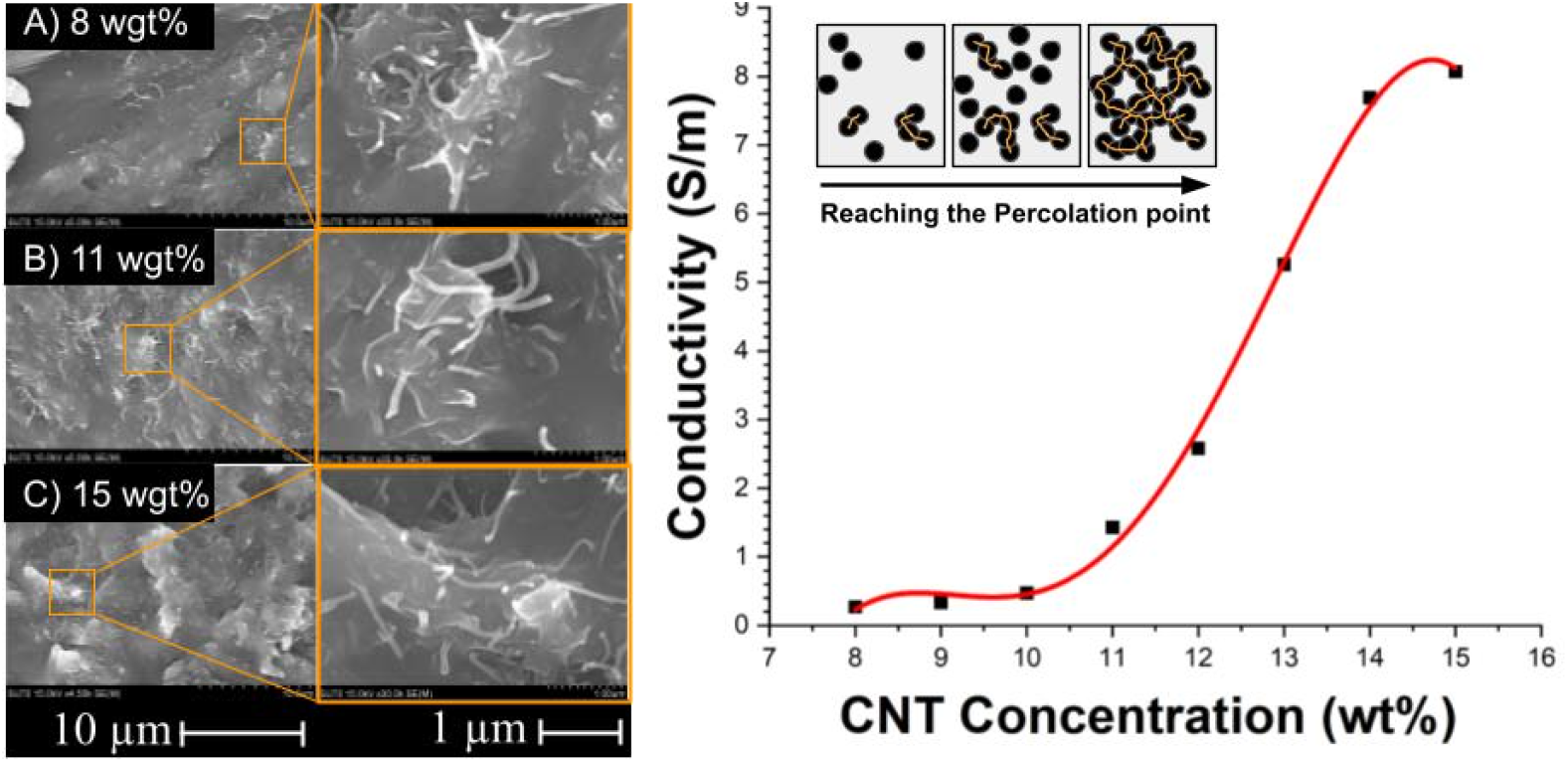
(A-C) SEM images of composite at different weight percent. As the composite reaches the percolation point the CNT causes mound-like structures to appear in the composite, this is due to non-homogenous mechanical properties in the composite. (D) Conductivity values of the composite at different weight percent.

### 3.3 Finite element analysis simulations

After the STL models were imported into Sim4life [48] and the simulations were completed [49], the peak simulated e-field was then recorded for each of the four coils, at the three corresponding depths in the phantoms. **Fig**.**9** shows the heat map intensity of each inspection plane when stimulated by the permender v-tip core. The simulated results were more linear and predictably spaced compared to the measured results, which was to be expected, the experimental measurements are subject to real world variations in coil placement, and phantom conductivity that will always remain highly controlled in a simulation. The peak intensities at 1000A stimulator output for the permender v-tip core, permender flat core, AISI 1010 v-tip core, and the AISI 1010 flat tip core were 110 V/m, 85 V/m, 100V/m, and 84 V/m respectively. The permender v-tip had the highest intensity stimulation.

**Figure 9:**
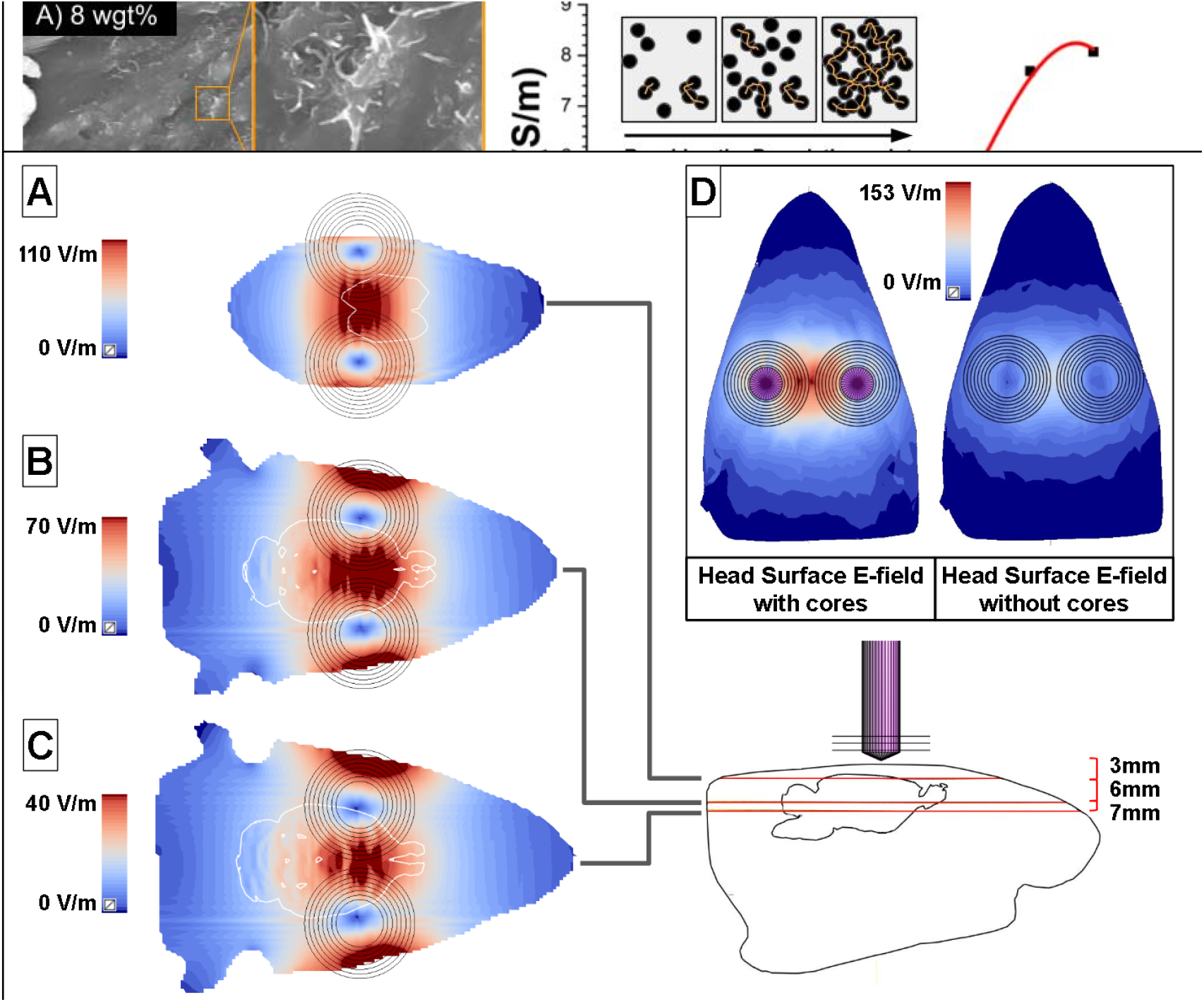
E-field distribution on the rat head model and planes, using the permendur v-tip coil configuration. (A) The three slices of interest that were discussed earlier are overlayed with the opacity lowered for visual effect. The scale of each of these slices is given on the left. (B) Sagittal and coronal view of the simulated results. In these images two scales were used for visual effect, the first scale describes the peak electric field of 158 V/m on the surface of the skin, the second scale describes the peak electric field of 100 V/m on the surface of the grey matter.

### 3.4 Measured e-fields in phantoms

Three phantoms were prepared and stimulated. The depth of each probe changed between phantoms, the first being at the brain surface 3mm from the coil, next was 6mm from the coil, and the was 7mm from the coil. Since we used a modified commercial stimulator was used on small handmade coils, we only went up to 15% of the stimulators maximum output to not damage the coils. The data presented in **Figure 10** shows the depth of the dipole probes and the focality of each individual coil. **Figure 10** also describes the core geometries and color codes the core material. The second posterior TDP in each phantom produced erroneous results due to connectivity and placement issues, therfore these measurements were not considered in our results. The core that produced the highest intensity stimulation of 115 V/m was the permender v-tip core. Both the AISI 1010 v-tip and the permender flat tip had very similar e-field strength at 94.6 V/m and 91.9 V/m respectively, showing that the higher permeability of the permender core gave higher field strengths even at less optimal core geometreis. The coil that produced the lowest cortical stimulation had the AISI 1010 flat-tip core, with an E-field of 85.9 V/m. The cortical probe had the most consistent results, while the mid-brain probe showed some nonlinearity.

**Figure 10.**
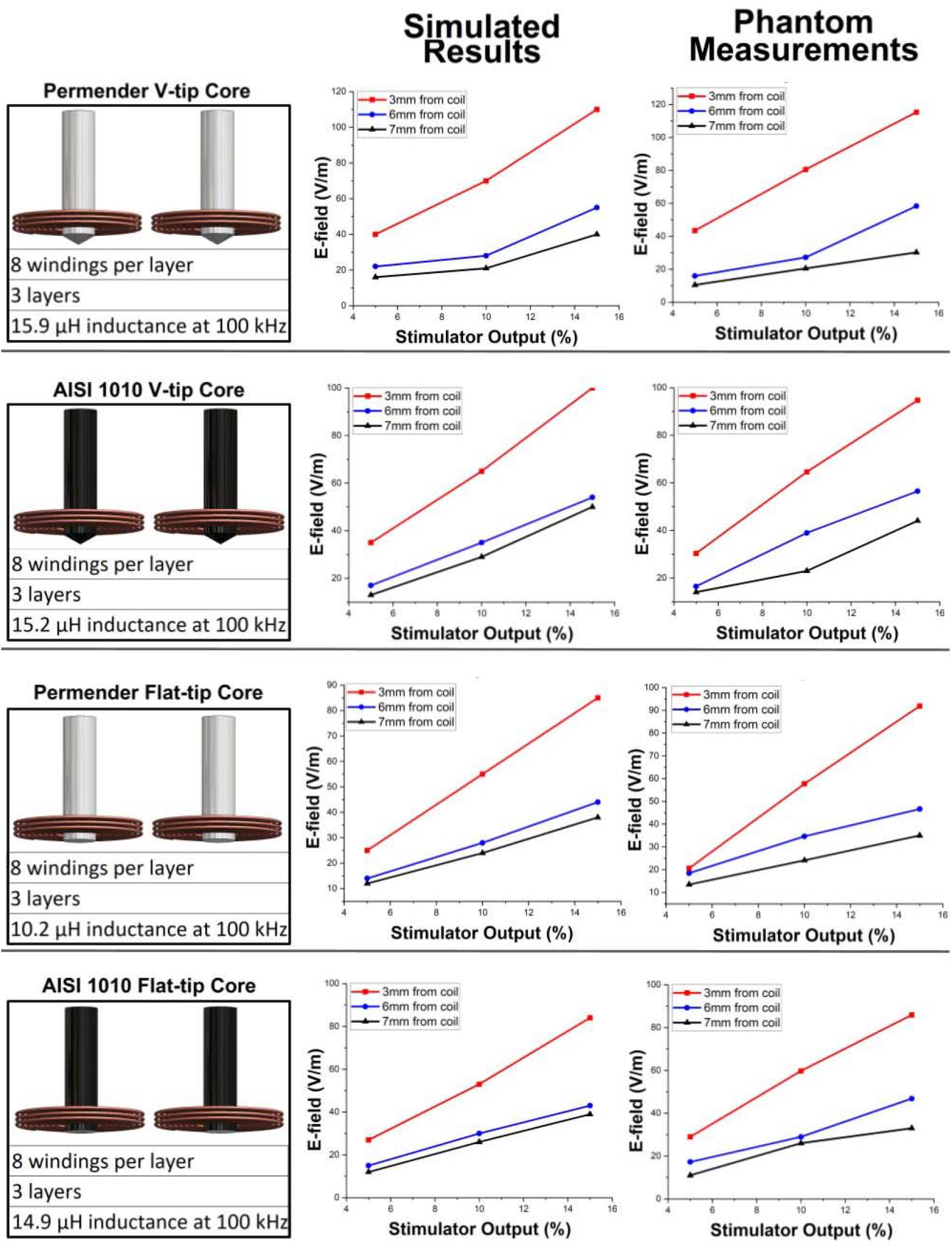
Simulated and measured e-field values plots using the 4 different coil configurations and the 3 phantoms. The values of the e-field exhibit similar trends for each coil and phantom

## 4. Discussion

The measured and simulated E-field were similar. The slight discrepancy between them could be due to several different factors including differences in the positioning of the coil, differences between the conductivity of the layers in the simulated model and the layers in the phantom, and differences in the STL models versus the physical phantoms and coils. Particularly the former may have a large impact on the results, while all the layers in the phantom were nearly identical to the STL files, the STL files of the coils were an approximation of the actual coil, because they were wound by hand. Although the simulated and mesured results were not exactly the same, the measured results were still relevant because they represent a good estimation of what the E-field values may be in actual rat models. This will allow for a quick and easy assessment of new coils and TMS protocols, especially in the cortical regions of the brain, before moving to *in vivo* settings. The cortical probe measurements followed the same trends as the simulations suggested, though the two subcortical probes did not exactly follow the same trends noted in the simulations. This difference could be attributed to a higher focality in the V tip shaped cores. Since the hotspot in these cores was smaller, precise targeting is more impactful, especially at lower depths. The phantoms measured the E-field vector of a 3mm x 3mm volume and did not capture the entire field at all points.

Moving forward, we suggest potential avenues for improvement particularly in creating separate gray and white matters with distinct conductivities. While this study validates the use of anatomical rat TMS phantom, the end result is not a phantom that would be useful to clinicians and those not familiar with the instrumentation used here. For a clinically useful TMS phantom, a user interface friendly to the less technical would be required, though this is easily solved. Our individualized rat brain phantoms offer a potentially valuable tool for TMS researchers and practitioners, bridging the gap between theoretical simulations and practical applications for those in the neuromodulation and TMS field.

## Conclusion

By combining MRI and CT data with image segmentation, 3D printing, and injection molding techniques, we created an anatomically detailed model that captures key aspects of rat cranial and brain structure. The use of a CNT-PDMS composite enabled tunable electrical conductivity to match biological tissues, while embedded triaxial dipole probes allowed in situ measurement of E-field vectors across multiple depths. Experimental measurements across different coil geometries and core materials revealed trends in field strength and focality that were generally consistent with finite element simulations, particularly for cortical measurements. The permender v-tip core produced the most intense and focal stimulation, highlighting the influence of both material properties and coil design on E-field distribution. While minor discrepancies between simulation and measurement were observed, likely due to geometric and conductivity differences, the phantom successfully enabled a controlled and repeatable environment for evaluating TMS devices. This work provides a scalable framework for developing similar phantoms for other animal models or even human studies and offers further evidence for increasing field focality and intensity using ferromagnetic cores.

## Acknowledgements

Authors would like to acknowledge the following NSF grants: # 2304513, #2349694 and #2336233. This project was also partially funded by VCU Presidential QUEST grant. P. S and C. D were funded by the NIH Brain Initiative 1R01NS112183 and the NIH Center for Mesoscale Mapping P41EB030006.

## Data Availability

Data from this study is available on the Open Science Framework at https://osf.io/aqtwh/

## References

[1] D. J. Stultz, S. Osburn, T. Burns, S. Pawlowska-Wajswol, and R. Walton, “Transcranial Magnetic Stimulation (TMS) Safety with Respect to Seizures: A Literature Review,” Neuropsychiatr Dis Treat, vol. 16, pp. 2989–3000, Dec. 2020, doi: 10.2147/NDT.S276635.

[2] M. Kobayashi and A. Pascual-Leone, “Transcranial magnetic stimulation in neurology,” Lancet Neurol, vol. 2, no. 3, pp. 145–156, Mar. 2003, doi: 10.1016/s1474-4422(03)00321-1.

[3] O. of the Commissioner, “FDA permits marketing of transcranial magnetic stimulation for treatment of obsessive compulsive disorder,” FDA. Accessed: Mar. 07, 2023. [Online]. Available: https://www.fda.gov/news-events/press-announcements/fda-permits-marketing-transcranial-magnetic-stimulation-treatment-obsessive-compulsive-disorder

[4] F Wang et al., “Low-Intensity Focused Ultrasound Stimulation Ameliorates Working Memory Dysfunctions in Vascular Dementia Rats via Improving Neuronal Environment,” Front Aging Neurosci, vol. 14, p. 814560, Feb. 2022, doi: 10.3389/fnagi.2022.814560.

[5] T. Dufor, A. M. Lohof, and R. M. Sherrard, “Magnetic Stimulation as a Therapeutic Approach for Brain Modulation and Repair: Underlying Molecular and Cellular Mechanisms,” Int J Mol Sci, vol. 24, no. 22, p. 16456, Nov. 2023, doi: 10.3390/ijms242216456.

[6] Y. Kozorovitskiy, R. Peixoto, W. Wang, A. Saunders, and B. L. Sabatini, “Neuromodulation of excitatory synaptogenesis in striatal development,” eLife, vol. 4, p. e10111, Nov. 2015, doi: 10.7554/eLife.10111.

[7] X. Niu, K. Yu, and B. He, “Transcranial Focused Ultrasound Induces Sustained Synaptic Plasticity in Rat Hippocampus,” Brain Stimul, vol. 15, no. 2, pp. 352–359, 2022, doi: 10.1016/j.brs.2022.01.015.

[8] M. S. George et al., “Daily Left Prefrontal Transcranial Magnetic Stimulation Therapy for Major Depressive Disorder: A Sham-Controlled Randomized Trial,” Archives of General Psychiatry, vol. 67, no. 5, pp. 507–516, May 2010, doi: 10.1001/archgenpsychiatry.2010.46.

[9] S. George et al., “Daily repetitive transcranial magnetic stimulation (rTMS) improves mood in depression,” Neuroreport, vol. 6, no. 14, pp. 1853–1856, Oct. 1995, doi: 10.1097/00001756-199510020-00008.

[10] R. B. Lipton and S. H. Pearlman, “Transcranial magnetic simulation in the treatment of migraine,” Neurotherapeutics, vol. 7, no. 2, pp. 204–212, Apr. 2010, doi: 10.1016/j.nurt.2010.03.002.

[11] L. Dinur-Klein et al., “Smoking cessation induced by deep repetitive transcranial magnetic stimulation of the prefrontal and insular cortices: a prospective, randomized controlled trial,” Biol Psychiatry, vol. 76, no. 9, pp. 742–749, Nov. 2014, doi: 10.1016/j.biopsych.2014.05.020.

[12] B. V. Watts, B. Landon, A. Groft, and Y. Young-Xu, “A sham controlled study of repetitive transcranial magnetic stimulation for posttraumatic stress disorder,” Brain Stimul, vol. 5, no. 1, pp. 38–43, Jan. 2012, doi: 10.1016/j.brs.2011.02.002.

[13] P. Cirillo et al., “Transcranial magnetic stimulation in anxiety and trauma-related disorders: A systematic review and meta-analysis,” Brain Behav, vol. 9, no. 6, p. e01284, Jun. 2019, doi: 10.1002/brb3.1284.

[14] M.-J. Ahmadizadeh and M. Rezaei, “Unilateral right and bilateral dorsolateral prefrontal cortex transcranial magnetic stimulation in treatment post-traumatic stress disorder: A randomized controlled study,” Brain Res Bull, vol. 140, pp. 334–340, Jun. 2018, doi: 10.1016/j.brainresbull.2018.06.001.

[15] J. D. Rollnik et al., “High frequency repetitive transcranial magnetic stimulation (rTMS) of the dorsolateral prefrontal cortex in schizophrenic patients,” Neuroreport, vol. 11, no. 18, pp. 4013–4015, Dec. 2000, doi: 10.1097/00001756-200012180-00022.

[16] S. S. Shin et al., “Transcranial magnetic stimulation and environmental enrichment enhances cortical excitability and functional outcomes after traumatic brain injury,” Brain Stimulation, vol. 11, no. 6, pp. 1306–1313, Nov. 2018, doi: 10.1016/j.brs.2018.07.050.

[17] A. A. Abdelrahman, M. Noaman, M. Fawzy, A. Moheb, A. A. Karim, and E. M. Khedr, “A double-blind randomized clinical trial of high frequency rTMS over the DLPFC on nicotine dependence, anxiety and depression,” Sci Rep, vol. 11, no. 1, p. 1640, Jan. 2021, doi: 10.1038/s41598-020-80927-5.

[18] Z.-D. Deng, S. H. Lisanby, and A. V. Peterchev, “Electric field depth–focality tradeoff in transcranial magnetic stimulation: Simulation comparison of 50 coil designs,” Brain Stimulation, vol. 6, no. 1, pp. 1–13, Jan. 2013, doi: 10.1016/j.brs.2012.02.005.

[19] M. Tashli, A. Mhaskar, G. Weistroffer, M. S. Baron, and R. L. Hadimani, “Novel multi-magnetic material transcranial magnetic stimulation coils for small animals application,” AIP Advances, vol. 14, no. 1, p. 015324, Jan. 2024, doi: 10.1063/9.0000772.

[20] Y. Meng, R. L. Hadimani, L. J. Crowther, Z. Xu, J. Qu, and D. C. Jiles, “Deep brain transcranial magnetic stimulation using variable ‘Halo coil’ system,” Journal of Applied Physics, vol. 117, no. 17, p. 17B305, Mar. 2015, doi: 10.1063/1.4913937.

[21] L. Liu et al., “Design and evaluation of a rodent-specific focal transcranial magnetic stimulation coil with the custom shielding application in rats,” Front Neurosci, vol. 17, p. 1129590, Apr. 2023, doi: 10.3389/fnins.2023.1129590.

[22] A. K. Kiani et al., “Ethical considerations regarding animal experimentation,” J Prev Med Hyg, vol. 63, no. 2 Suppl 3, pp. E255–E266, Oct. 2022, doi: 10.15167/2421-4248/jpmh2022.63.2S3.2768.

[23] L. J. Crowther, R. L. Hadimani, A. G. Kanthasamy, and D. C. Jiles, “Transcranial magnetic stimulation of mouse brain using high-resolution anatomical models,” Journal of Applied Physics, vol. 115, no. 17, p. 17B303, Jan. 2014, doi: 10.1063/1.4862217.

[24] H. Magsood and R. L. Hadimani, “Development of anatomically accurate brain phantom for experimental validation of stimulation strengths during TMS,” Materials Science and Engineering: C, vol. 120, p. 111705, Jan. 2021, doi: 10.1016/j.msec.2020.111705.

[25] H. Magsood, F. Syeda, K. Holloway, I. C. Carmona, and R. L. Hadimani, “Safety Study of Combination Treatment: Deep Brain Stimulation and Transcranial Magnetic Stimulation,” Front. Hum. Neurosci., vol. 14, Apr. 2020, doi: 10.3389/fnhum.2020.00123.

[26] “NITRC: Templates for Long-Evans Rat Brain: Tool/Resource Info.” Accessed: Dec. 02, 2024. [Online]. Available: https://www.nitrc.org/projects/tpm_rat/

[27] “NITRC: SIGMA Rat Brain Templates and Atlases: Tool/Resource Info.” Accessed: Dec. 07, 2024. [Online]. Available: https://www.nitrc.org/projects/sigma_template

[28] “Quintuple AISI 1010 carbon steel core coil for highly focused transcranial magnetic stimulation in small animals,” ResearchGate. Accessed: Nov. 21, 2024. [Online]. Available: https://www.researchgate.net/publication/348959483_Quintuple_AISI_1010_carbon_steel_core_coil_for_highly_focused_transcranial_magnetic_stimulation_in_small_animals

[29] M. Tashli, G. Weistroffer, A. Mhaskar, D. Kumbhare, M. S. Baron, and R. L. Hadimani, “Investigation of soft magnetic material cores in transcranial magnetic stimulation coils and the effect of changing core shapes on the induced electric field in small animals,” AIP Advances, vol. 13, no. 2, p. 025319, Feb. 2023, doi: 10.1063/9.0000550.

[30] R. Casanova et al., “Biological parametric mapping: A statistical toolbox for multimodality brain image analysis,” NeuroImage, vol. 34, no. 1, pp. 137–143, Jan. 2007, doi: 10.1016/j.neuroimage.2006.09.011.

[31] S. J. Kiebel, J. Ashburner, J.-B. Poline, and K. J. Friston, “MRI and PET Coregistration—A Cross Validation of Statistical Parametric Mapping and Automated Image Registration,” NeuroImage, vol. 5, no. 4, pp. 271–279, May 1997, doi: 10.1006/nimg.1997.0265.

[32] D. Arnone et al., “Computational meta-analysis of statistical parametric maps in major depression,” Human Brain Mapping, vol. 37, no. 4, pp. 1393–1404, 2016, doi: 10.1002/hbm.23108.

[33] N. J. Kazemi et al., “Ictal SPECT statistical parametric mapping in temporal lobe epilepsy surgery,” Neurology, vol. 74, no. 1, pp. 70–76, Jan. 2010, doi: 10.1212/WNL.0b013e3181c7da20.

[34] V. Tavares, D. Prata, and H. A. Ferreira, “Comparing SPM12 and CAT12 segmentation pipelines: a brain tissue volume-based age and Alzheimer’s disease study,” Journal of Neuroscience Methods, vol. 334, p. 108565, Mar. 2020, doi: 10.1016/j.jneumeth.2019.108565.

[35] E. R. Sowell, P. M. Thompson, C. J. Holmes, R. Batth, T. L. Jernigan, and A. W. Toga, “Localizing Age-Related Changes in Brain Structure between Childhood and Adolescence Using Statistical Parametric Mapping,” NeuroImage, vol. 9, no. 6, pp. 587–597, Jun. 1999, doi: 10.1006/nimg.1999.0436.

[36] “Development of Anatomically Accurate Brain Model of Small Animals for Experimental Verification of Transcranial Magnetic Stimulation | IEEE Journals & Magazine | IEEE Xplore.” Accessed: Dec. 02, 2024. [Online]. Available: https://ieeexplore.ieee.org/abstract/document/9512033

[37] A. Fedorov et al., “3D Slicer as an Image Computing Platform for the Quantitative Imaging Network,” Magn Reson Imaging, vol. 30, no. 9, pp. 1323–1341, Nov. 2012, doi: 10.1016/j.mri.2012.05.001.

[38] Y.-N. Chen et al., “Stereotaxic atlas of the infant rat brain at postnatal days 7–13,” Front. Neuroanat., vol. 16, Aug. 2022, doi: 10.3389/fnana.2022.968320.

[39] P. M. Glover and R. Bowtell, “Measurement of electric fields due to time-varying magnetic field gradients using dipole probes,” Phys Med Biol, vol. 52, no. 17, pp. 5119–5130, Sep. 2007, doi: 10.1088/0031-9155/52/17/001.

[40] A. Gaikwad, M. Olowe, and S. Desai, “Deformation Mechanism of Aluminum, Copper, and Gold in Nanoimprint Lithography Using Molecular Dynamics Simulation,” Nanomaterials (Basel), vol. 13, no. 24, p. 3104, Dec. 2023, doi: 10.3390/nano13243104.

[41] S. Liu, H. Yoshioka, A. Kuwahata, and M. Sekino, “Optimizing transcranial magnetic stimulator coils for minimal influence of individual variability in head geometry,” AIP Advances, vol. 14, no. 2, p. 025325, Feb. 2024, doi: 10.1063/9.0000795.

[42] L. I. Navarro de Lara et al., “A 3-axis coil design for multichannel TMS arrays,” NeuroImage, vol. 224, p. 117355, Jan. 2021, doi: 10.1016/j.neuroimage.2020.117355.

[43] P. P. C. Bhagubai and J. F. P. Fernandes, “Multi-Objective Optimization of Electrical Machine Magnetic Core Using a Vanadium–Cobalt–Iron Alloy,” IEEE Transactions on Magnetics, vol. 56, no. 2, pp. 1–9, Feb. 2020, doi: 10.1109/TMAG.2019.2950880.

[44] Y. Fu et al., “Fe–Co-based crystalline soft magnetic coatings with ultra-high saturation magnetization above 1.9T via co-axial powder feeding plasma-transferred arc welding,” J Mater Sci: Mater Electron, vol. 34, no. 6, p. 468, Feb. 2023, doi: 10.1007/s10854-023-09831-8.

[45] “Dielectric Properties□» IT’IS Foundation.” Accessed: Dec. 09, 2024. [Online]. Available: https://itis.swiss/virtual-population/tissue-properties/database/dielectric-properties/

[46] A. S. Asan, S. Gok, and M. Sahin, “Electrical fields induced inside the rat brain with skin, skull, and dural placements of the current injection electrode,” PLoS ONE, vol. 14, no. 1, p. e0203727, Jan. 2019, doi: 10.1371/journal.pone.0203727.

[47] M. Sekino, H. Ohsaki, S. Yamaguchi-Sekino, N. Iriguchi, and S. Ueno, “Low-frequency conductivity tensor of rat brain tissues inferred from diffusion MRI,” Bioelectromagnetics, vol. 30, no. 6, pp. 489–499, 2009, doi: 10.1002/bem.20505.

[48] L. J. Crowther, R. L. Hadimani, and D. C. Jiles, “Effect of Anatomical Brain Development on Induced Electric Fields During Transcranial Magnetic Stimulation,” IEEE Transactions on Magnetics, vol. 50, no. 11, pp. 1–4, Nov. 2014, doi: 10.1109/TMAG.2014.2326819.

